# Cannabinoid CB2R regulates T cell gut-homing in preclinical Crohn’s model

**DOI:** 10.1101/2025.09.05.674422

**Authors:** Robert S. Leddy, Cinthia L. Hudacheck, Hannah M. Phelan, Billy P. Egan, Carol M. Aherne, Julián Romero, Cecilia J Hillard, Paul Jedlicka, Colm B. Collins

**Affiliations:** School of Biomolecular and Biomedical Science, University College Dublin, Dublin, Ireland; UCD Conway Institute of Biomolecular and Biomedical Research, University College Dublin, Dublin, Ireland; Department of Medicine, Division of Gastroenterology, Hepatology & Nutrition, University of Colorado, Anschutz Medical Campus; School of Medicine, University College Dublin, Dublin, Ireland; Faculty of Experimental Sciences, Universidad Francisco de Vitoria, Pozuelo de Alarcón, Spain; Department of Pharmacology and Toxicology, Neuroscience Research Center, Medical College of Wisconsin, Milwaukee, WI, USA; Department of Pathology, University of Colorado, Anschutz Medical Campus, Colorado USA

## Abstract

Leukocyte trafficking is a critical step in development of chronic intestinal diseases such as Crohn’s disease. While strategies that block gut-homing have yielded partial success, this disease remains uncurable leaving an unmet clinical need. This is the first paper to describe a role for cannabinoid receptor two (CB_2_R) signalling in promoting retinoic acid-mediated induction of the gut-homing associated integrin heterodimer α4β7. Using *in vitro* and *in vivo* models, we characterised the effects of pharmacological CB_2_R agonists and inverse agonists on T cell homing receptor expression and transmigration across gut-associated endothelial barriers. This ERK-dependent process coincides with increased T cell adherence in response to CB_2_R agonism with JWH133. These effects were reversed with an inverse agonist GP-1a in a CB_2_R dependent manner. Selective deletion of CB_2_R using CRISPR in vitro or CD4^Cre/+^ floxed mice in vivo resulted in impaired endothelial cell adherence and decreased diapedesis into the ileal lamina propria. T cell-specific deletion of *cnr2*, the gene encoding CB_2_R, attenuated chronic murine ileitis characterised by decreased naïve T cell infiltration and loss of tissue architecture in 20wk TNF^ΔARE/+^mice. This study supports further therapeutic development of CB_2_R-blocking drugs for the treatment of inflammatory bowel disease.

## Introduction

Inflammatory bowel disease (IBD), encompassing Crohn’s disease (CD) and ulcerative colitis (UC), is a chronic, relapsing inflammatory disorder of the gastrointestinal tract characterised by dysregulated immune responses and impaired intestinal homeostasis. People with IBD may experience severe chronic visceral pain which is challenging to manage, in part because of the negative impact of opioid analgesics on digestive function and epithelial repair ^1, 2^.

Cannabis is seen by patients as a potential means to alleviate chronic pain and motility dysfunction associated with IBD^3^. The endocannabinoid system (ECS) signals primarily via two G protein-coupled receptors, CB_1_R and CB_2_R. These are G protein-coupled receptors that primarily couple to Gi/o proteins^4^, leading to inhibition of adenylate cyclase, decreased cyclic adenosine monophosphate (cAMP) levels, and modulation of downstream signalling pathways such as MAPKs (ERK1/2, p38, JNK)^5–7^. These pathways regulate diverse cellular processes including cytokine production, cell migration, proliferation, and apoptosis. This is further regulated by their distinct expression patterns and functions. CB_1_R, abundant in the central nervous system, regulates neurotransmission including the psychoactive effects synonymous with cannabis products^8^, while also regulating intestinal secretion and motility via expression on enteric neurons^9^. Conversely, CB_2_R while certainly induced in the brain during inflammation^10, 11^, is predominantly expressed in immune tissues (spleen, lymph nodes, bone marrow) and hematopoietic cells, including T cells, macrophages, and neutrophils, at levels up to 100-fold higher than CB_1_R. This immune-focused distribution identifies CB_2_R as a potential regulator of inflammation and, lacking psychoactive effects, highlights its appeal as a therapeutic target for treating inflammatory diseases.

In the context of intestinal inflammation, both animal models and human studies have demonstrated upregulation of CB_2_R and increased levels of ECS ligands in the gut^12, 13^. For instance, CB_2_R expression is markedly increased on infiltrating immune cells in experimental colitis models^14, 15^ and in the inflamed mucosa of IBD patients^12, 16–18^. Similarly, elevated levels of ECS ligands anandamide and 2-AG have been observed in the intestinal tissues and plasma of individuals with active IBD, correlating with disease severity^19–21^.

Furthermore, selective cannabinoid receptor two (CB_2_R) agonist, APD371, has proven effective in clinical trials at decreasing visceral pain^22^. However, clinical trials with phytocannabinoids consistently fail to demonstrate efficacy in disease modification. A randomised study of Crohn’s disease (CD) patients revealed symptomatic improvement without achieving increased clinical remission, while a trial in ulcerative colitis (UC) showed no improvement in remission rates between cannabis and placebo groups despite quality-of-life benefits^23, 24^. A 2021 trial further corroborated the dissociation between subjective relief and objective inflammatory markers^25^, a finding reinforced by a Cochrane meta-analysis highlighting insufficient evidence for cannabis in IBD remission induction or maintenance^26^. Of greater concern is that prolonged cannabis use (>6 months) correlates with worsened CD outcomes, including a five-fold increase in surgical risk^27^, underscoring unresolved controversies regarding its therapeutic role.

CB_2_R also appears to play a role in regulating immune cell trafficking, a process critical for maintaining intestinal immune surveillance and homeostasis. In the experimental autoimmune encephalomyelitis mouse model of multiple sclerosis, CB_2_R activation has been demonstrated to have protective effects by reducing CD4^+^ T cell infiltration and microglial activation, leading to improved disease outcomes^28^. Conversely, CB_2_R-deficient mice exhibited worse disease outcomes, including increased CD4^+^ T cell infiltration of the brain, highlighting the receptor’s importance in this regard^28^. Similarly, chronic administration of tetrahydrocannabinol (THC), an active compound found in *cannabis*, to simian immunodeficiency virus (SIV)-infected male rhesus monkey increased intestinal integrin α4β7^+^ CD4^+^ T cells, CD8^+^ T cells, and central memory T cells and the significant increase in the expression of pro-inflammatory cytokines within the gut^29^. Our lab has previously demonstrated that inverse agonism of CB_2_R attenuates inflammation in a preclinical Crohn’s model with a concomitant decrease in infiltrating T cells in the murine ileal lamina propria^12^. Dysregulation of leukocyte trafficking contributes to the chronic inflammation characteristic of IBD and may contributed to cannabinoid-mediated exacerbation of chronic disease. Gut-specific homing is primarily driven by α4β7 expression on circulating T cells interacting with mucosal addresssin cell adhesion molecule MAdCAM-1 on endothelial cells^30^.

Retinoic acid (RA) induces stable α4β7 expression on CD4^+^ T cells via epigenetic regulation, ensuring that even when these cells migrate away from the required RA signal, they retain the imprinted gut-homing “zip-code” resulting in long-lasting gut-homing T cells. During priming, RA blocks the methylation of specific CpG sites upstream of the *ITGA4* promoter in mice and humans, leading to the stable expression of α4β7 but not α4β1^31^. Interestingly, CB_2_R activation has been shown to mimic these epigenetic events induced by RA via the triggering the same histone modifications^32^, meaning it may be possible that receptor activation may promote gut-homing properties. Alternatively, CB_2_R may regulate integrin expression via MAPK. CB_2_R’s role in MAPK signalling was first identified by Bouaboula *et al.* who showed that CB_2_R activation leads to ERK1/2 phosphorylation^33^. Subsequent studies have found CB_2_R activation to trigger p38 MAPK in lymphoma cells and induce the activation of ERK1/2 and JNK MAPK in microglial cells^34^. CB_2_R activation has also been noted to lead to the phosphorylation of ERK1/2 and subsequent proinflammatory cytokine release in a human epithelial cells epithelial cells ^35^. Activation of CB_2_R also induces transient phosphorylation of ERK1/2 in T cells ^36^.

In psoriasis, integrin expression correlates with MAPK activation, implicating MAPK activation in disease pathogenesis^37^. Although this does not directly implicate MAPK signalling as a possible mechanism in CB_2_R-mediated integrin expression, the manner in which CB_2_R signalling alters MAPK signalling provides a plausible explanation as to how CB_2_R signalling alters the expression of various adhesion molecules in multiple cells types.

In addition to its effects on immune cell trafficking, CB_2_R signalling modulates cytokine production and immune cell differentiation, contributing to the resolution of inflammation and tissue repair. Notably, CB_2_R activation suppresses the production of proinflammatory cytokines and promotes the expansion of regulatory T cell populations, which are often deficient in IBD. These immunomodulatory properties position CB_2_R as a promising therapeutic target for the treatment of IBD and other inflammatory disorders. Despite these insights, the precise role of CB_2_R and the ECS in IBD pathogenesis remains incompletely understood. The upregulation of CB_2_R and ECS ligand levels in inflamed tissues suggests a compensatory, protective response, yet clinical outcomes with exogenous cannabinoids are inconsistent at best. It is possible that chronic cannabis use may desensitize or dysregulate the ECS, undermining its potential benefits. Furthermore, the complex interplay between CB_2_R signalling, adhesion molecule expression, and immune cell trafficking warrants further investigation to elucidate the mechanisms underlying cannabis effects in IBD.

## Methods

### Cellular Adhesion Assays

Jurkat T cells (2.5×10⁶ cells/mL) were cultured in RPMI-1640 and treated in quadruplicate for 48 h with vehicle treatment, 1 µM retinoic acid (RA) ± 1 µM JWH-133 or 10 µM GP-1a. Cells were counted, resuspended in HBSS + 0.5 mM MnCl₂, and added to 24-well plates pre-coated with recombinant MAdCAM-1 (3 ng/µL, Bio-Techne #6056-MC). After 30 min at 37°C, non-adherent cells were removed, and adhered cells were trypsinised (200 µL, 2–3 min, RT; Gibco #25300-054) and recounted. Adhesion percentage was calculated as (adhered cells / total cells) × 100.

### MAdCAM-1-Dependent Cellular Adhesion Assay

Jurkat T cells (2.5×10⁶ cells/mL in RPMI-1640) were treated in quadruplicate with 1 µM retinoic acid (RA) ± 1 µM JWH-133 (CB2R agonist) or 10 µM GP-1a (CB2R inverse agonist) for 48 h; vehicle controls received media alone. Concurrently, HEK293T cells (ATCC, Manassas, VA, USA) stably expressing MAdCAM-1 (1×10⁶ cells/well in DMEM supplemented with 10% FBS, 1% penicillin-streptomycin, and 1.24% L-glutamine (ThermoFisher) were seeded to achieve >90% confluency after 48 h. Jurkat cells were harvested, stained with 5 µM carboxyfluorescein diacetate (CFDA; ThermoFisher, V12883) in PBS (15 min, 37°C), and washed. CFDA-labelled Jurkat cells (1 mL) were added to MAdCAM-1-expressing HEK293T monolayers and co-cultured for 30 min at 37°C. Non-adherent cells were aspirated, and adhered Jurkat cells were detached with 200 µL trypsin (2–3 min, RT; Gibco #25300-054). Adherent cells were quantified via fluorescence microscopy (Olympus CKX41/EP50) and expressed as before.

For cellular adhesion assays under shear stress, m-Slide I Luer 3D slides (Ibidi, #87176) and supplemented DMEM were degassed overnight at 37°C. On the following day, slides were coated with 3.3 µg/mL Rat Collagen I (Cultrex, #3440-100-01) in PBS for 2 h at 37°C, followed by application of a collagen gel matrix composed of 60% collagen, 10% 10X PBS (ThermoFisher, #70013-016), 14% NaHCO₃, 16% DMEM, and 46% deionised water, with polymerisation for 2 h at 37°C. Human umbilical vein endothelial cells (HUVECs, ATCC) were maintained in supplemented DMEM, labelled in serum-free DMEM at 2×10⁶ cells/mL with 1 µM CellTracker Orange CMTMR (Invitrogen, #C2927) for 30 min at 37°C, washed, and seeded into the slides at 1.6×10⁶ cells/mL, then cultured overnight. Wild-type and CNR2⁻/⁻ Jurkat T cells were resuspended at 3×10⁶ cells/mL and treated with 1 mM JWH-133 in T25 flasks (Greiner, #C6356) for 24 h at 37°C. The following day, Jurkat cells were stained with 0.33 µM CellTracker Green CMFDA (Invitrogen, #C7025) for 30 min at 37°C, washed, and resuspended in pre-warmed, degassed, phenol red-free RPMI (ThermoFisher, #11835063) at 0.1×10⁶ cells/mL. Cell suspensions were loaded into syringe pumps and perfused through the HUVEC-coated slides, which had been preconditioned with phenol red-free RPMI, at a shear stress of 0.4 dyn/cm² for 18 h (flow rate calculated per manufacturer’s instructions). After perfusion, slides were washed with PBS, fixed with 4% paraformaldehyde for 30 min at room temperature, and washed again prior to imaging on an Olympus FV1000 confocal microscope. Adhered Jurkat cells were quantified from three images per chamber, with the mean value used for analysis.

### Jurkat T Cell Gene Expression Analysis

Jurkat E6-1 human T lymphocytic leukaemia cell line, from the American Type Culture Collection (ATCC, Manassas, Virginia, USA) were maintained at 37°C (95% O_2_/5% CO_2_) in RPMI-1640 containing L-glutamine (ThermoFisher Scientific) supplemented with 1% penicillin-streptomycin (ThermoFisher Scientific) and 10% fetal bovine serum. Jurkat T cells plated at 2.5x10^6^ cells/ml were treated for 48hr with Retinoic Acid (RA; 1µM; Merck; Cat No. R2625), with(out) CB_2_R agonist JWH-133 (1µM; Tocris BioScience) or CB_2_R inverse agonist GP-1a (10µM; Tocris BioScience) or vehicle as appropriate. Cells were harvested, washed in PBS and total RNA extracted using the RNeasy Mini kit (Qiagen) according to the manufacturer’s instructions. Eluted RNA samples were then sent for transcriptomic profiling sequencing by BGI Genomics, with an RNA integrity number of ≥ 6.5 used as RNA-quality cut-off for inclusion in analysis. Raw sequencing data was converted to FASTQ file format using Illumina and uploaded to Galaxy, before quality control of reads was done using the Cutadapt pipeline. The cleaned reads were then mapped to a reference genome using STAR and checked using the Integrative Genomics Viewer. From the mapped sequences, the number of reads per annotated genes were counted using the featurecounts package. The DESeq2 package was then used on the read counts to normalise and extract differentially expressed genes (DEGs) between treatment groups. Functional enrichment and pathway analysis of the DEGs was performed using the Fast Gene Set Enrichment Analysis (fgsea) package to extract any interesting gene ontologies. The full analysis dataset is available from https://www.ncbi.nlm.nih.gov/bioproject/PRJNA1301825. For expression analysis, isolated RNA was transcribed using a high-capacity cDNA Reverse Transcription Kit (Applied Biosystems). Relative quantification of mRNA expression was performed using Taqman Gene Expression Assays and the QuantStudio 12K Flex Real-Time PCR System. Real-time PCR (RT-PCR) assays for ITGB7 (ThermoFisher Scientific; Hs01565750_m1), ITGA4 (ThermoFisher Scientific; Hs00168433_m1), ITGAE (ThermoFisher Scientific; Hs01025372_m1), ITGB1 (ThermoFisher Scientific; Hs01127536_m1) and CNR2 (ThermoFisher Scientific; Hs05019229_s1) were carried out, with ribosomal 18s (Applied Biosciences; Cat No. 4319413E) as an endogenous control.

### Biological Network Analysis

Biological network, functional, and pathway analyses were performed using Ingenuity Pathway Analysis (IPA, Ingenuity Systems, Redwood City, CA, USA). For canonical pathway and network analyses, gene lists containing up- and down-regulated transcripts (with corresponding log2 fold change and adjusted p-values) from RNA sequencing of CB2R activation (JWH-133) or blockade (GP-1a) conditions were imported into IPA. No stringent filtering criteria were applied, enabling comprehensive identification of transcriptionally regulated pathways and networks. IPA determined the degree of pathway regulation based on log2 fold change and –log10(adjusted p-value) for each gene.

### Dual Luciferase Promoter Reporter Assays

Jurkat T cells were seeded in 24-well plates (2.5×10⁵ cells/well) and transfected with 1 µg DNA/well using Lipofectamine™ LTX with PLUS™ Reagent (ThermoFisher). Constructs included full-length or truncated ITGB7 sequences cloned into pGL4 vectors (GenScript), co-transfected with Renilla luciferase plasmid (9:1 ratio). Cells were treated 24 h post-transfection as indicated. Lysates resuspended in 100 µL Passive Lysis Buffer (Promega Dual-Luciferase® Assay) were analysed for firefly and Renilla luciferase activities using a ClarioStar® PLUS plate reader (BMG LabTech). Normalised relative light units (RLUs) were calculated and visualised in GraphPad Prism 10.

### Mice

TNF^ΔARE/+^ mice (B6.129S-Tnftm2GKI/Jarn; MGI:3720980) were previously generated by backcrossing heterozygous TNF^ΔARE/+^ mice to C57BL/6J. CD4Cre and RAG1^−/−^ mice (Jackson Laboratory) were bred in-house and crossed with CB_2_R^Fl/Fl^ mice^38^ (provided by C.J. Hillard and J. Romero Paredes) to generate CB_2_R^Fl/Fl^CD4^Cre^TNF^ΔARE/+^ triple transgenics; Cre-negative littermates served as controls.

### Flow cytometry and staining

For cytokine profiling, splenocytes, mesenteric lymphocytes, and lamina propria mononuclear cells were stimulated for 3 h at 37°C with 50 ng/mL PMA (Sigma #P8139), 1 µg/mL ionomycin (Sigma #I0634), and 10 µg/mL brefeldin A (Sigma #B7651) in RPMI-1640 (10% FBS, 2 mM L-glutamine, 1% penicillin-streptomycin). Cells were permeabilized and stained with antibodies against CD62L (MEL-14, BioLegend), CD4 (GK1), CD45 (30-F11), CD44 (IM7), IL-10 (JES5-16E3), IL-17A (TC11-18H10.1), IL-17F (9D3.1C8), IFN-γ (XMG1.2), and FoxP3 (MK-14; all BioLegend), preceded by Fc block (BioLegend #93). Live cells were identified using Live/Dead Fixable Aqua dye (Invitrogen). Samples fixed in 1% PFA were analysed on a FACSCanto II (BD Biosciences) with FlowJo v10.8.1. Histological assessments were performed by a blinded pathologist as described previously.

### Phosphorylation Status Assessment by Proteome Profiler Antibody Array

Phosphorylation profiles of intracellular signalling components following CB2R modulation were analysed using the Human Phospho-Kinase Array™ (R&D Systems, ARY003C). Jurkat T cells (2.5×10⁶ cells/mL) were treated for 10 min in RPMI-1640 containing 1 mM retinoic acid ± 1 mM CB2R agonist JWH-133. Lysates normalized via BCA assay (ThermoFisher) were processed using 300 μg protein per assay membranes (A/B) per manufacturer protocol. Signal intensities were quantified with ImageJ (v1.53).

### Assessing Intracellular/Extracellular Target Expression via Flow Cytometry

Jurkat T cells were seeded at 2.5×10⁶ cells/mL in RPMI-1640 and treated in triplicate with 1 µM retinoic acid (RA) with or without 1 µM JWH-133 for 48 hours at 37°C; vehicle controls received media alone. Following incubation, cells were transferred to V-bottom plates, centrifuged at 1500 × g for 5 minutes, and resuspended in Hank’s Balanced Salt Solution (HBSS) with or without 0.5 mM manganese chloride for integrin activation. For surface staining, cells were incubated with Alexa Fluor 647-conjugated anti-human integrin α4β7 antibody (Bio-Techne/R&D Systems, FAB10078R) for 30 minutes on ice prior to fixation with 4% paraformaldehyde at 4°C. For intracellular staining, cells were fixed and permeabilized using a commercial buffer set, then stained with Alexa Fluor 647-conjugated anti-human integrin α4β7 (Bio-Techne/R&D Systems, FAB10078R), APC-conjugated anti-phospho-ERK1/2 (T202/Y204) (ThermoFisher Scientific, 17-9109-42), PE-conjugated anti-phospho-LCK (Y394) (Bio-Techne/Novus Biologicals, NBP3-13305), or eFluor 660-conjugated anti-phospho-LCK (Y505) (ThermoFisher Scientific, 50-9076-42) as appropriate. Live cells were identified using Live/Dead Fixable Aqua dye. Following staining, cells were resuspended in PBS and analysed on a CytoFLEX LX flow cytometer (Beckman Coulter), with data processed using CytExpert software.

### Statistics

Statistical analyses were performed using a two-tailed t test. Values of p < 0.05 were statistically significant. Graphs show mean +/- standard error of the mean (SEM) and were generated using GraphPad Prism software. Values of P<0.05 were considered statistically significant

## Results

### Increased T cell adherence in response to pharmacological CB_2_R activation

To assess whether cannabinoid-treatment would make Jurkat T cells more capable of binding to the gut-specific adhesion molecule MAdCAM-1, we utilised static cellular adhesion assays utilising MAdCAM-1 coated plates (**Figure 1A**) and MAdCAM-1-expressing HEK293T cells (**Figure 1B and C**). Retinoic acid (1µM) was also added to induce expression of gut-homing integrins on the T cell surface. Consistent with the trend observed at a promoter and transcriptional, CB_2_R inverse agonism with GP-1a (10 µM) significantly reduced the adherence of treated Jurkat T cells to MAdCAM-1 coated plated and HEK293T endothelial-like cells in both assays. Conversely, CB_2_R activation with JWH133 (1 µM) led to a significant increase in the percentage of adhered cells in both assays following 48 h cannabinoid treatment. In a more biologically relevant manner, we also assessed the ability of cannabinoid-treated WT and *CNR2*^-/-^ Jurkat T cells to bind to MAdCAM-1-expressing HUVECs under shear flow. Here, the treated T cells were circulated through a fluidics system at a flow rate of 0.4 dyn/cm^3^ over a static HUVEC endothelial monolayer, with the percentage of adhered, fluorescently tagged T cells to the endothelial monolayer being recorded via use of fluorescent confocal microscopy. Consistent with the static assays used, this assay also highlighted the role of CB_2_R signalling in mediating functional T cell adhesion. The role of CB_2_R activation in inducing the expression of functional α4β7 and mediating adhesion to MAdCAM-1 was observed to be receptor-dependent in this assay, with receptor knockdown mediating a significant decrease in the relative percentage of adhered cells when compared to WT cells (**Figure 1D & 1E**). Finally, we examined the contribution of CB_2_R to leukocyte trafficking *in vivo* using a competitive homing assay. Isolated fluorescently labelled CD4^+^ T cells from WT and CB_2_R^Fl/Fl^CD4^Cre+^ splenocytes were injected i.p. into RAG1^-/-^ lymphopenic hosts and expression of relative abundance of each genotype assed after 24 h. CD4^+^ T cells from mice with T cell-specific deletion of CB_2_R displayed a significant decrease in frequency in the ileal lamina propria and draining mesenteric lymph node (MLN), while levels of splenic cells were increased. Colonic cell numbers were unaffected which may reflect distinct means of gut homing (**Figure 1F**). Taken together, this data indicates the role of CB_2_R signalling in regulating T cell gut-homing both *in vitro* and *in vivo*.

**Figure 1.**
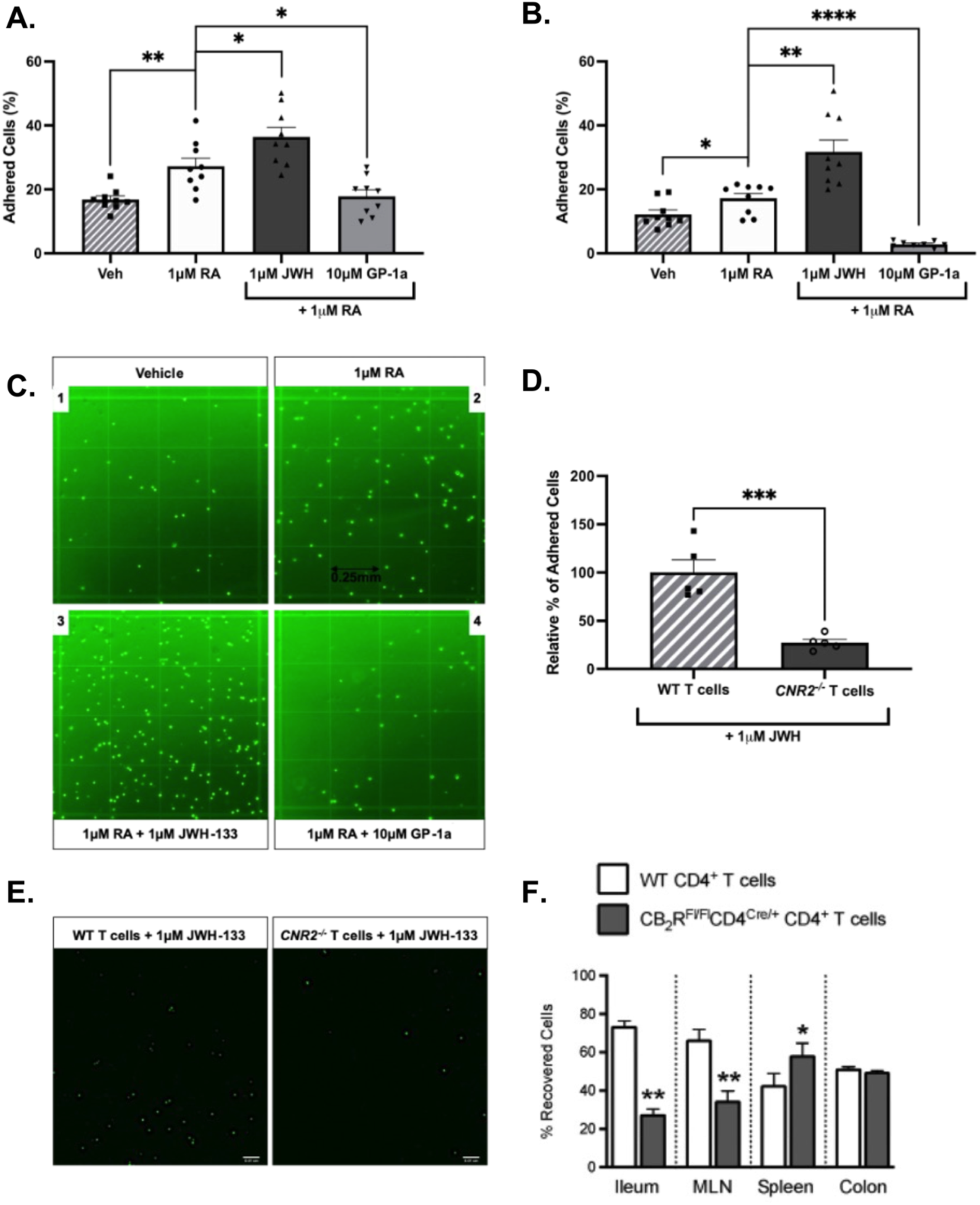
CB_2_R mediated regulation of integrin β7 protein expression. CB_2_R signalling induces an increase in integrin β7 protein expression that functionally binds to MAdCAM-1, in a receptor-dependent manner. Static adhesion assays assessing the ability of the CB_2_R-mediated β7 to functionally bind to **(A)** MAdCAM-1-coated plates and **(B)** and **(C)** MAdCAM-1-expressing HEK293T cells. **(D)** Cellular adhesion assay under shear flow using fluorescently-tagged WT and CNR2^-/-^ Jurkat T cells to assess the requirement of CB_2_R in inducing the expression of functional β7 that binds to MAdCAM-1-expressing HUVECs. Results represent mean±SEM for n=3 independent experiments. * p<0.05, **p< 0.01, *** p<0.001, ****P<0.0001.

### CB_2_R-mediated regulation of T cell transcription

To better understand how CB2R signalling might drive an increase in gut-homing, we performed bulk RNA sequencing on Jurkat T cells in the presence of the required RA signal. RNA was sequenced from Jurkat T cells treated with 24 h CB_2_R blockade and activation, as well as vehicle-treated controls, identifying distinct transcriptional profiles of genes regulated by the receptor. Respectively, 241 and 142 differentially expressed genes were highlighted following receptor blockade or activation (**Figure 2A&B**), and a number of these were significantly altered in expression based on their log2 fold-change when compared to vehicle-treated cells (**Figure 2C&D**). These gene-sets were obtained via the digital removal of RA-regulated genes, generating two lists made up of genes specifically regulated following CB_2_R blockade or activation (**Figure 2E**). Using these distinct sets of differentially expressed genes, pathway analysis was used to get a broad view of the specific pathways and gene interaction networks differentially regulated following manipulation of CB_2_R signalling. Several pathways pivotal in normal T cell functioning were highlighted, including cellular immune responses and cellular metabolism (**Supplemental Figure S1 & S2**), indicating the broad role of the receptor in regulating multiple facets of normal T cell functioning. In the context of the cellular immune response pathways identified, genes such as *CCR4, ITGB7,* and *ITGAL* were identified as being some of the top differentially regulated genes (**Figure 2C & D**), whereas *MYC, PIK3R6,* and *HKDC1* were amongst the top 50 differentially expressed genes following CB_2_R manipulation in the context of cellular metabolism, each playing a key role in glucose metabolism specifically. This RNA sequencing data and process of gene-set generation was validated using RT-qPCR, with the mRNA expression of several selected candidate genes of interest being shown to mirror the RNA sequencing data in a treatment-specific manner (not shown). Pathway analysis also revealed various in-depth gene-gene interaction networks following receptor blockade and activation, allowing for a broad view of the potential genetic interactions that may be driving CB_2_R-mediated responses in T cells. Taken together, CB_2_R signalling drives transcriptional alterations within T cells, with receptor blockade and activation yielding two distinctly different transcriptional profiles that appear to be important in several normal T cell function including cellular immune responses and glucose metabolism.

**Figure 2.**
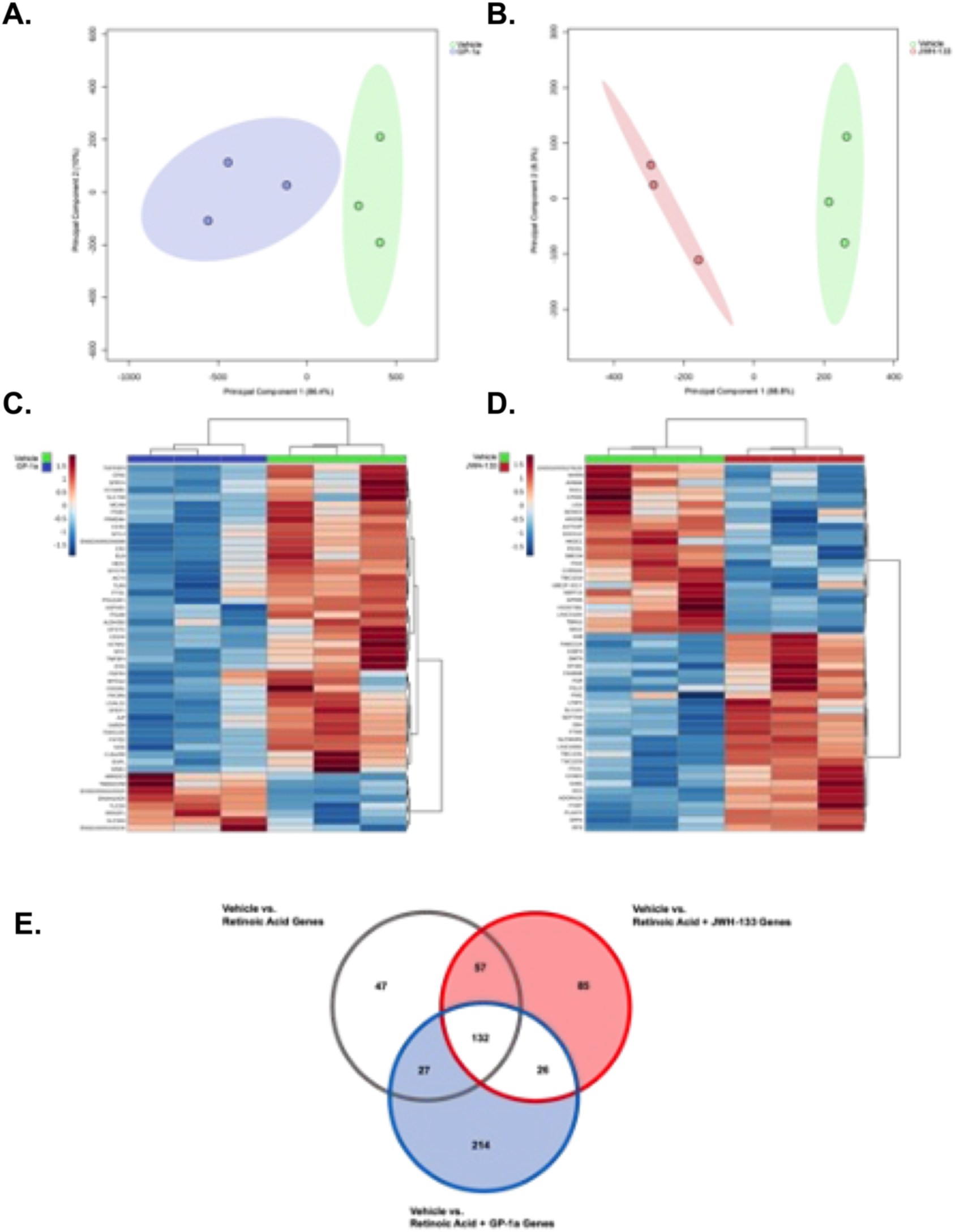
CB_2_R drives distinct transcriptional profiles in Jurkat T cells. Bulk RNA sequencing highlighting distinct transcriptional profiles following CB_2_R modulation (N = 3). Principal component analysis (PCA) score plots for the genes regulated following CB_2_R blockade with GP-1a (**A**) or (**B**) activation with JWH133. R2 = 0.903 and R2 = 0.87, respectively. (**C**, **D**) Heatmaps of the top 50 genes based on Log2FoldChange specifically regulated following CB_2_R blockade or activation, respectively. The degree of expression is indicated by +1.5 (red) and -1.5 (blue). (**E**) Venn diagram of the degree of overlap of differentially expressed genes between treatment groups indicative of the level of change induced by CB_2_R signalling.

### CB_2_R-mediated regulation of T cell integrin β7 mRNA and protein expression

Given the impact of CB_2_R on driving gut homing and the differential expression of integrin β7 identified by RNAseq, we next examined potential regulation of integrin β7 by CB_2_R at a transcriptional and protein level. Jurkat T cells were transfected with an integrin β7 dual luciferase reporter assays to assess β7 promoter activity in response to cannabinoid treatment in the presence of RA as before. As previously described RA alone significantly increased β7 promoter activity. Following 24 h and 48 h cannabinoid treatment, we observed a significant increase in reporter activity with CB_2_R activation and a corresponding decrease in response to receptor blockade (**Figure 3A**). Transcriptional regulation of CB_2_R signalling on integrin β7 was also validated using RT-PCR, indicating integrin β7 mRNA expression to be significantly downregulated and upregulated following CB_2_R blockade and activation, respectively (**Figure 3B**). As with the dual luciferase reporter studies, an earlier timepoint of 6 h was also investigated, but resulted in no differences amongst treatment groups (not shown). For scientific rigour and to assess whether the effect of CB_2_R signalling on integrin expression was β7-specific, integrins α4, αE, and β1 mRNA expression levels were also investigated, but no significant alterations in their expression were found across all timepoints tested. This indicates that CB_2_R signalling regulates integrin expression on T cells in a manner that is selective for integrin β7.

**Figure 3.**
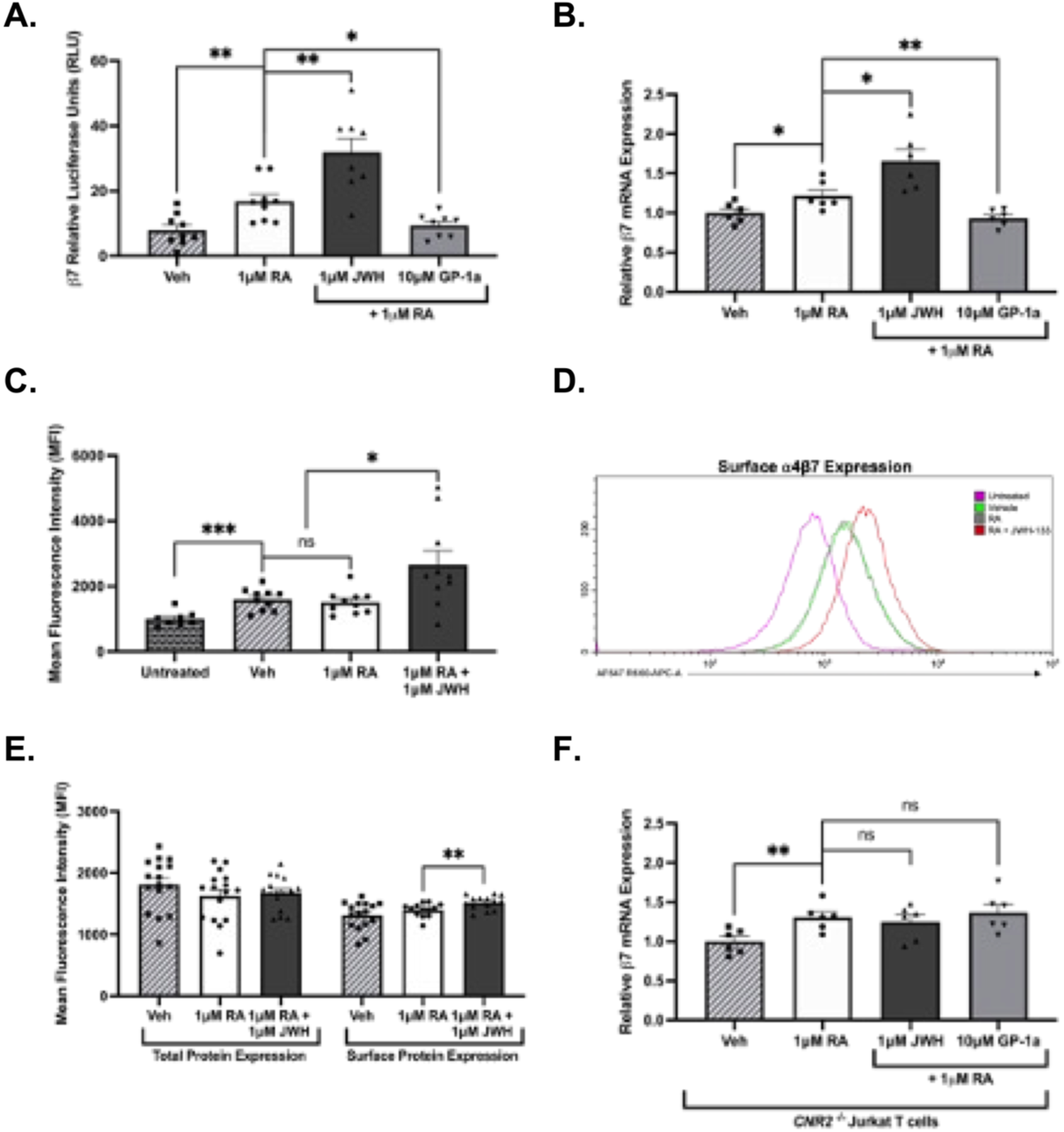
CB_2_R activation drives surface expression of integrin α4β7. **(A)** Integrin β7 promoter activity assessed using dual luciferase reporter assay after 24 h cannabinoid treatment in the presence of 1μM retinoic acid. **(B)** Integrin β7 mRNA expression levels assessed using RT-PCR following 24 h cannabinoid treatment. **(C)** Flow cytometry analysis of integrin expression levels at a surface protein level by means of mean fluorescence intensity (MFI) following 48 h CB_2_R activation in Jurkat T cells. (**D**) Representative histogram overlay of the average MFI values for each treatment group, displaying alterations in surface α4β7 expression on T cells following CB_2_R activation. (**E**) Flow cytometry analysis used to make comparisons between total and surface integrin α4β7 protein expression levels in Jurkat T cells after 48 h CB_2_R activation. (**F**) RT-PCR studies investigating integrin β7 mRNA expression levels in CRISPRi CNR2^-/-^ Jurkat T cells following 24 h cannabinoid treatment. Results represent mean±SEM for n≥3 independent experiments.* p<0.05, ** p<0.01, *** p<0.001.

Whilst interesting to note the role of CB_2_R in regulating the expression of integrin β7 at both promoter and transcriptional levels, it was also important to assess whether these upstream alterations translated to changes in α4β7 surface expression at a protein level. This was assessed using flow cytometry. To induce a conformational change in integrin α4β7 expression towards an activated isoform to allow for effective integrin-antibody binding, the cells were pretreated with manganese chloride (MnCl_2_), which induces a conformational change in integrin presentation facilitating labelling antibody binding. Following 48 h CB_2_R activation, integrin β7 protein expression was significantly elevated on the surface of T cells, consistent with increased translation of upregulated *ITGB7* mRNA, as indicated by MFI (**Figure 3C and D**). The requirement for MnCl_2_ pretreatment was confirmed via flow cytometry analysis without the presence of MnCl_2_ which failed in facilitating effective integrin-antibody binding and therefore resulted in no changes between any treatment groups at that timepoint (not shown). We then assessed both the surface α4β7 protein expression and the whole-cell total α4β7 protein expression in non-permeabilised and permeabilised cells, respectively. Interestingly, the elevated expression of α4β7 following CB_2_R activation was induced at a surface protein level, rather than in a total protein manner (**Figure 3E**).

Following the transfection of a *CNR2* CRISPRi complex under the control of a tetracycline-inducible promoter, we were able to deduce that the significant modulation of integrin β7 expression in T cells following CB2R manipulation occurs in a receptor-dependent manner, with those cells with inducible *CNR2* knockout failing to have any sort of alterations in β7 expression following 24 h cannabinoid treatment (**Figure 3F**). Additionally, *CNR2* knockout had no effect on RA-induced integrin β7 expression. Taken together, this data highlights the expansive role of CB_2_R signalling in regulating the expression of the gut-homing integrin β7 on T cells at both a promoter and transcriptional level, in a manner that is receptor-dependent.

### Ras/Raf/MEK/ERK signalling is central to CB_2_R-mediated induction of integrin β7

To explore the mechanism involved in the regulation of CB_2_R-mediated integrin β7 expression on T cells, we used a β7 promoter truncation assay to aid in identifying the specific sequence responsible for CB_2_R-mediated β7 upregulation. Three β7 pGL4 plasmid vectors were transfected Jurkat T cells prior to cannabinoid treatment; Full Length (FL; ∼ 1000 base pairs), Truncation 1 (T1; ∼ 750 base pairs; **Figure 4A**), and Truncation 2 (T2; ∼ 500 base pairs; **Figure 4B**). CB_2_R activation significantly increased β7 promoter activity relative to RA-treated samples following 48 h cannabinoid treatment in those cells transfected with T1 and T2. In contrast, those cells transfected with T2 failed to show any enhancement in integrin β7 promoter activity relative to RA-treated cells following CB_2_R activation, indicating the specific sequence for CB_2_R-mediated β7 expression to be located within the 250 base pairs between T1 and T2 (**Figure 4C**). Using the Alggen Promo database and the FASTA sequence of these 250 base pairs, we were able to map the specific transcription factor binding sites located along the sequence, with Ras/Raf/MEK/ERK pathway-associated transcription factors being the most prominent along the specific sequence (**Figure 4D**). To further validate whether Ras/Raf/MEK/ERK signalling was involved in the mechanism driving integrin β7 expression on T cells following CB_2_R activation, we utilised the pan-Raf inhibitor LY3009120. Dual luciferase reporter assays found this pan-Raf inhibitor to significantly block the CB_2_R-mediated induction in β7 expression in Jurkat T cells, whilst having no impact on RA-mediated increases, highlighting the role for this pathway in regulating CB_2_R-induced β7 expression specifically (**Figure 4E and F**). To summarise, this data reveals the specific genetic sequence responsible for CB_2_R-mediated integrin β7 expression on T cells, something that has not been described previously. Using this sequence, the Raf/Ras/MEK/ERK pathway, particularly ERK1/2 signalling, was found to be central in regulating integrin β7 expression on T cells following CB_2_R activation.

**Figure 4.**
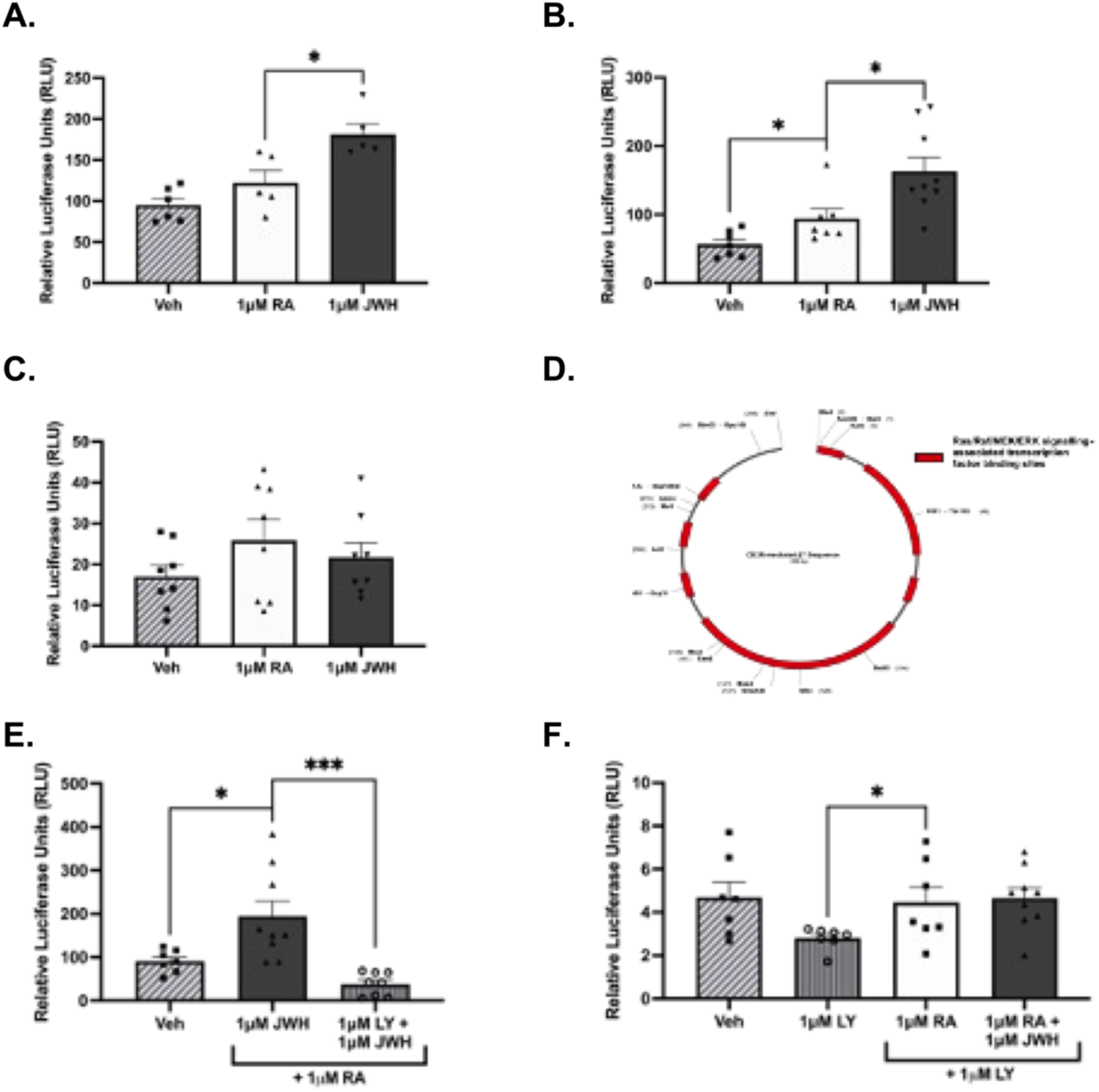
Transcription factor binding site mapping identifies Ras/Raf/MEK/ERK signalling as central to CB_2_R-mediated β7 induction. Truncations of the full length β7 promoter reporter revealed CB2R induced luciferase activity with Truncation 1 (T1; ∼ 750 base pairs; Figure 4A), and Truncation 2 (T2; ∼ 500 base pairs; Figure 4B) following 48 h CB_2_R activation. In contrast, cells transfected with T2 failed to show any enhancement in integrin β7 promoter activity relative to RA-treated cells following CB_2_R activation (Figure 4C), indicating the target sequence for CB_2_R-mediated β7 induction to be located within the 250 base pairs between T1 and T2. **(D)** Plasmid map schematic of the specific sequence responsible for CB_2_R-mediated β7 expression in Jurkat T cells. The location of Ras/Raf/MEK/ERK-associated transcription factor binding site locations are displayed in red. Schematic generated using SnapGene and mapped using the Alggen Promo database. **(E)** Dual luciferase reporter assay investigating the Ras/Raf/MEK/ERK-dependent nature of CB_2_R-mediated integrin β7 promoter activity in T cells, using a pan-Raf inhibitor. **(F)** Dual luciferase reporter assay highlighting how Ras/Raf/MEK/ERK inhibition targets the CB_2_R-mediated effects on integrin β7 promoter activity specifically, rather than that of RA. Results represent mean±SEM for n≥3 independent experiments.* p<0.05, ** p<0.01, *** p<0.001.

### Phosphokinases identify downstream targets of CB2R that may regulate integrin β7

To assess to what extent ERK1/2 signalling is involved in regulating integrin β7 expression on T cells following CB_2_R activation and to identify potential upstream or downstream regulators, a human phosphokinase dot blot array was used to profile differentially active protein kinases and to investigate the unique kinase network activated following receptor activation in T cells (**Figure 5A**). Of the 39 proteins tested in the array, members of the Src family of protein tyrosine kinases appeared to be altered following CB_2_R activation. These included Fgr, Lck, Lyn, Src, and Yes (**Figure 5B**). To explore what aspect of the Ras/Raf/MEK/ERK signalling pathway is central in regulating CB_2_R-mediated β7 expression, we focussed on ERK signalling due its downstream position in the pathway. A time-course flow cytometry analysis assay assessing the levels of phosphorylated ERK1/2 at the activation site at T202/Y204 following CB2R activation revealed significant increases in the phosphorylation levels of this protein following 30 minutes of cannabinoid treatment, and subsequent trends towards an increase after 45 and 60 minutes, albeit insignificant (**Figure 5C**). RA appeared to have no effects on the levels of ERK1/2 phosphorylation at any timepoint tested, again indicating the CB_2_R-specific nature of the involvement of this pathway in regulating integrin β7 expression on T cells (**Figure 5C**). To assess the role of Lck in this regard, a time-course flow cytometry analysis assay assessing the levels of phosphorylated Lck at the T394 activation site following CB_2_R activation was used. This assay revealed significant decreases in the phosphorylation levels of this kinase after 10 and 15 minutes following CB_2_R activation (**Fig. 5C**). Conversely, investigating the phosphorylation levels of Lck at the inhibitory site at T505 found significant increases after 15 and 30 minutes CB_2_R activation, further suggesting that Lck signalling is reduced following the activation of CB_2_R (**Figure 5D**). Again, RA treatment appeared to have no impact on Lck activity at any of the timepoints tested. Taken together, these findings suggest the Src family as being potentially important in the ERK signalling responsible for CB_2_R-mediated α4β7 expression on T cells, but in a manner that is independent of Lck kinase activity.

**Figure 5.**
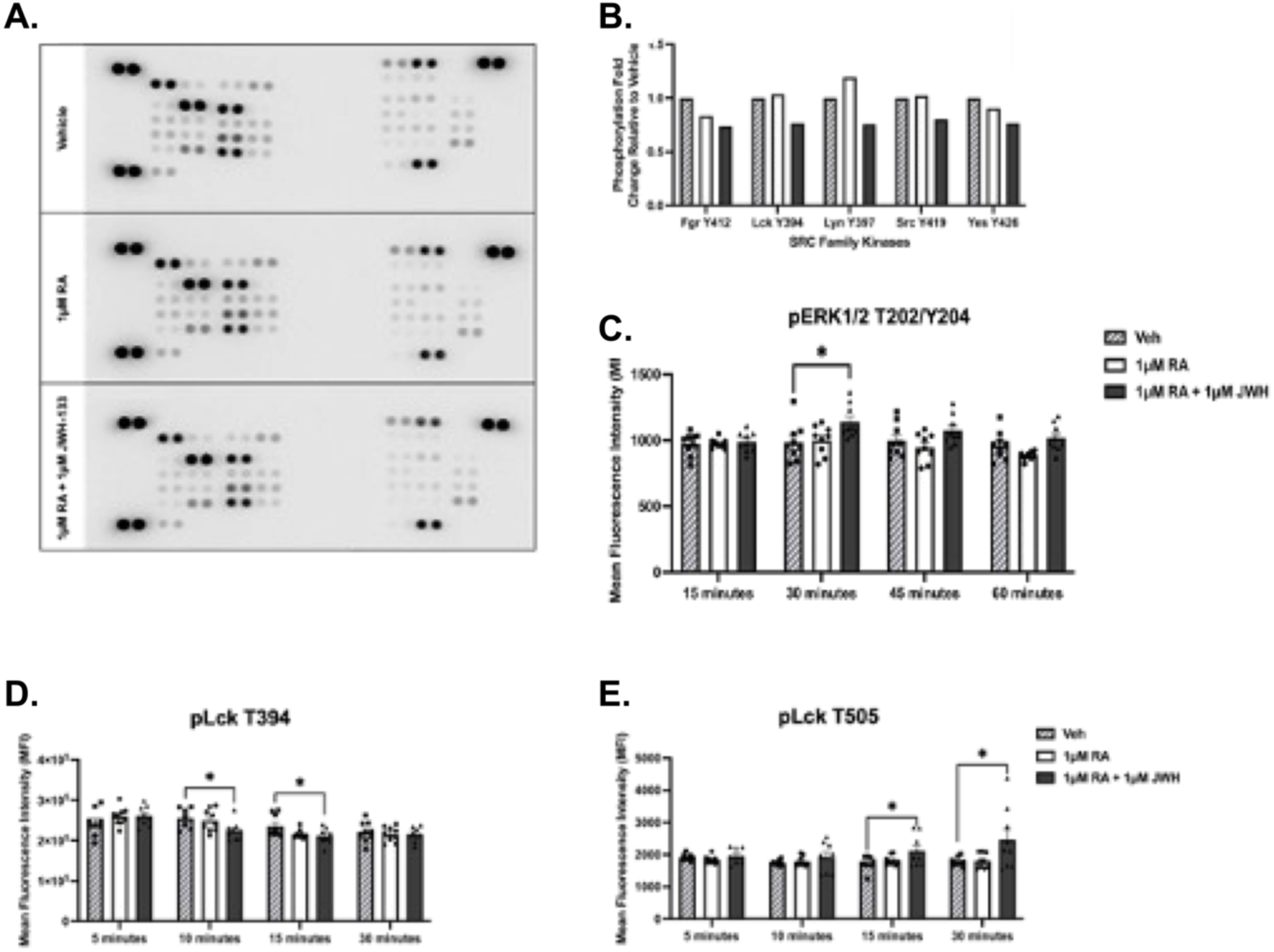
Phosphokinase array identifies potential downstream targets of CB_2_R that may mediate integrin β7 on T cells following receptor activation. **(A)** Phosphokinase array dot blot following 10 minute CB_2_R activation. Dot colour intensity is proportional to the degree of protein phosphorylation. **(B)** Densitometry highlighting Src family kinases as being differentially phosphorylated following CB_2_R activation. **(C)** Flow cytometry time-course assay assessing the levels of phosphorylated ERK1/2 at the activating site at T202/Y204 in response to CB_2_R activation. **(D)** Flow cytometry time-course assay assessing phosphorylated levels of Lck at the activating site T394 following CB_2_R activation. **(E)** Flow cytometry time-course assay assessing phosphorylated levels of Lck at the inhibitory site T505 following CB_2_R activation. Results represent mean±SEM for n≥3 independent experiments.* p<0.05.

### T cell-specific deletion of CB_2_R attenuates inflammation in TNF-driven chronic ileitis

Given the ability of CB_2_R signalling to promote T cell adherence and upregulation of gut=homing integrin β7, we next determined if T cell-mediated expression of CB_2_R would impact on disease pathogenesis in a chronic pre-clinical IBD model. Histological examination of inflammatory indices by a trained pathologist (PJ) blinded to the study, identified significant decreases in active inflammation, chronic inflammation, villus distortion and total inflammatory scores in 20-week-old TNF^βARE/+^CB_2_R^FL/FL^CD4^Cre/+^ compared to littermate controls (**Figure 6A & B**). This coincided with a significant decrease in inflammatory cell infiltrate in the spleen, MLN and ileal lamina propria (LP). Phenotypic analysis of the T cell subsets within the inflamed gut identified a concomitant increase in relative frequency of CD4^+^ FoxP3^+^ regulatory T cells (**Figure 6D**) as well as CD4^+^ IL-10^+^ anti-inflammatory T cells limited to the ileal LP (**Figure 6E**). Taken together this data supports the hypothesis that CB_2_R signalling exacerbates intestinal disease driving pro-inflammatory T cells into the inflamed intestine.

**Figure 6.**
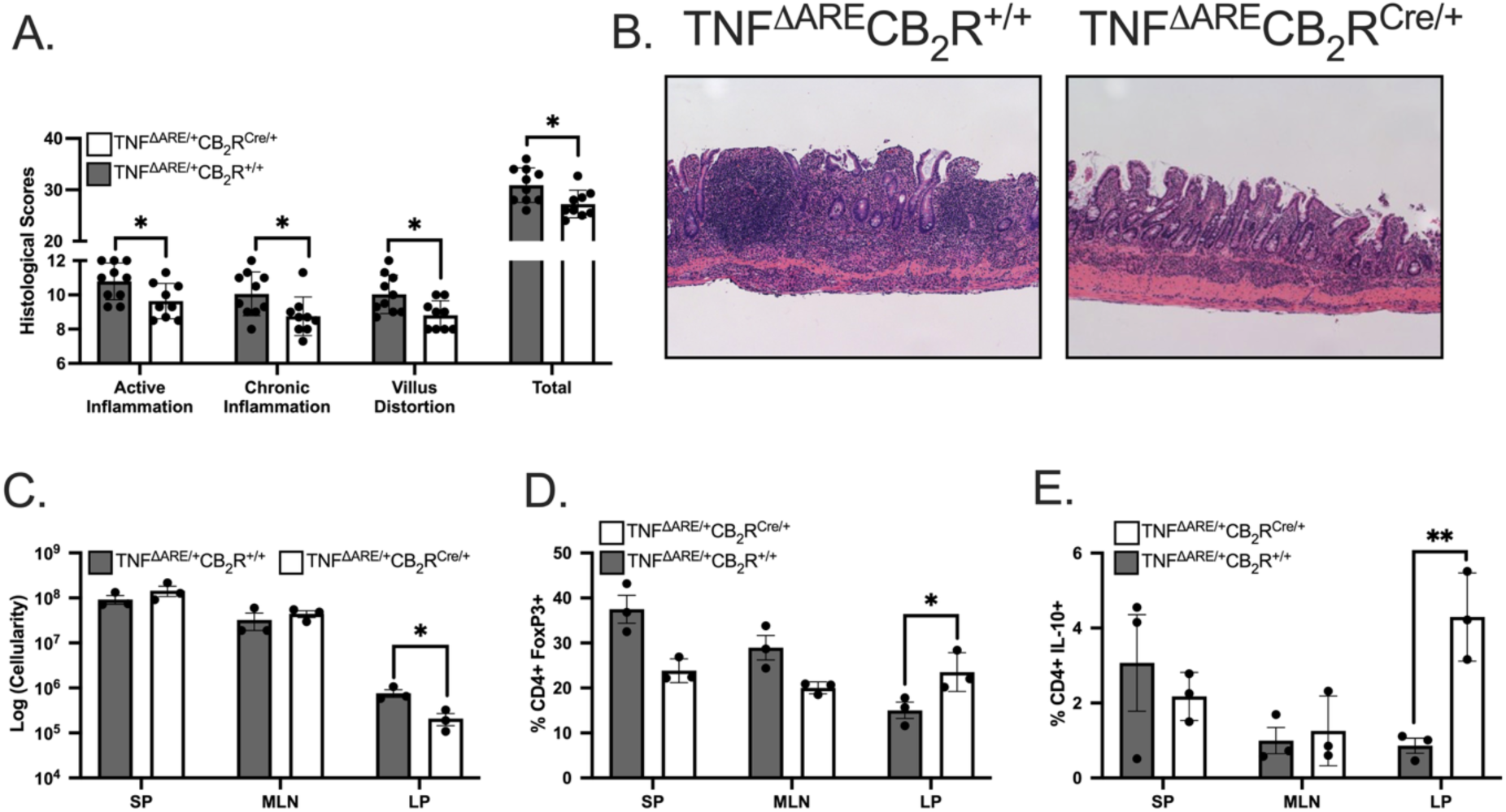
T cell-specific CB_2_R deletion attenuates chronic murine ileitis *in vivo.* (**A**) Blinded histological evaluation of intestinal inflammation in the 20wk TNF^ΔARE/+^CB^2^R^Fl/Fl^CD4^Cre/+^ ileum relative to cre controls demonstrated a significant decrease in acute and chronic inflammatory indices in addition to a reduction in villus distortion in TNF^ΔARE/+^ with selective deletion of CB_2_R from their T cells. (**B**) Representative micrographs indicating the improvement in villus architecture and reduction in inflammatory cell infiltrate in TNF^ΔARE/+^CB^2^R^Fl/Fl^CD4^Cre/+^ mice relative to controls. (**C**) Quantification of lamina propria lymphocytes recovered from indicated organs support the histological findings of a decrease in cell infiltration into the ileum. Flow cytometry identified a concomitant reduction in (**D**) CD4^+^ FoxP3^+^ regulatory T cells and (**E**) IL-10 producing CD4^+^ T the ileal lamina propria of 20-week-old mice. Results represent mean ± SEM for 4-7 mice per group from two or more independent studies. *P<0.05, **P<0.01.

### CB_2_R deletion coincides with a reduction central memory T cell expression

To better understand the subsets of T cells that are altered by deletion of CB2R signalling, we categorised live CD4^+^ T cells based on expression of CD44 and CD62L with cells expression high levels of CD62L and low CD44 expression referred to as naïve cells, CD62L^Low^CD44^High^ as effector cells and CD62L^High^CD44^High^ cells as central memory (**Figure 7A**). CB_2_R deletion drove a significant decrease in naïve and central memory T cell expression in T cells recovered from the ileal LP (**Figure 7B**) with those findings echoed in the draining MLN (**Figure 7C**). Whereas the number of naïve T cells were unaltered in the uninflamed spleen, levels of circulating central memory T cells were also significantly decreased in the spleen (**Figure 7D**) consistent with findings from the intestinal tissues. Taken together this points to a decrease in naïve T cell infiltration to the inflamed intestine resulting from a loss of CB_2_R signalling which we attribute to a reduced induction of integrin β7. Additionally, T cell-specific deletion of CB_2_R appears to also reduce central memory T cell phenotype which may be critical to the chronic nature of IBD.

**Figure 7.**
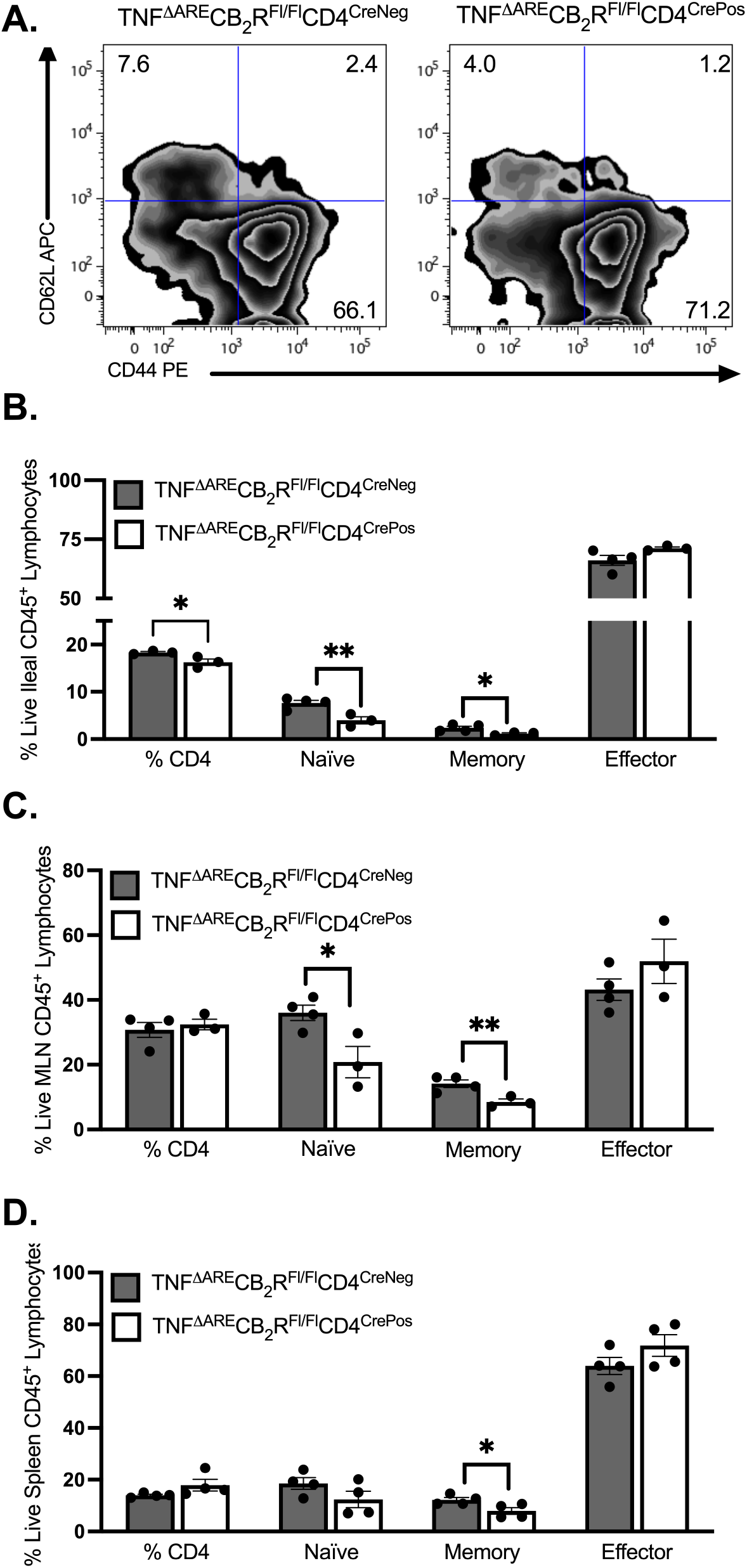
CB_2_R deficiency reduces ileal infiltration of naïve T cells *in vivo*. (**A**) Representative zebra plots showing flow cytometric analysis of the frequency of live CD4^+^ T cells expressing CD44 and CD62L from the inflamed ileum lamina propria (LP). Quantification of relative expression of CD4 T cells, CD44^Low^CD62L^High^ naïve T cells, CD44^High^CD62L^High^ central memory T cells and CD44^High^CD62L^Low^ effector T cells from the (**B**) ileum, (**C**) mesenteric lymph node (MLN) and (**D**) spleen (SP) demonstrate a significant decrease in CD4 T cells in the ileum only, a decrease in naïve T cell infiltration into both the LP and MLN along with a decrease in memory T cells in all three compartments tested, in 20-week-old TNF^ΔARE/+^ mice with selective deletion of CB_2_R from their T cells. Results represent mean ± SEM for 4 mice per group from two or more independent studies. *P<0.05, **P<0.01.

## Discussion

The goal of this study was to understand the contribution of T cell CB_2_R signalling to chronic intestinal inflammation. We had previously demonstrated that pharmacological CB_2_R inverse agonism with GP-1a attenuated murine ileitis but an argument could be made that the mechanism reflected targeting CB_2_R on other cell types^12^. T cell-specific CB_2_R deletion attenuated chronic murine ileitis and decreased leukocyte trafficking. Attempts to reproduce this *in vitro* failed unless RA signalling was also included. This led us to test the hypothesis that CB_2_R potentiates RA-mediated induction of gut-homing α4β7 on T cells leading to increased CD4^+^ T cell trafficking to the small intestine. This increase drives chronic murine ileitis *in vivo* in a model that is sensitive to disruption of leukocyte trafficking^39–41^. In the presence of RA, CB_2_R agonism upregulated integrin β7 at both message and protein level in both murine and human T cells. Selective CB_2_R activation on human leukocytes has also been shown to downregulate active forms of the α4β1 integrin complex^42^, adding further weight to this hypothesis. While prior studies have linked CB_2_R to the regulation of integrins such as α4β1, αLβ2, and αMβ2, our findings are the first to implicate CB_2_R in the modulation of α4β7 expression^43, 44^. Our results indicate that CB_2_R activation enhances RA-driven upregulation of α4β7. This synergistic interaction is consistent with previous reports in other systems, where CB_2_R signalling combines with suboptimal RA to promote bone growth and potentiate RAR-mediated transcription in hepatocytes^45, 46^. Mapping the sequence conferring CB_2_R-dependent regulation of β7 integrin expression on T cells using truncated promoter-reporters revealed multiple predicted RXR-α binding sites upstream of the β7 start site. RXR-α, a retinoid X receptor that heterodimerizes with RARs, likely mediates RA signalling at these loci, suggesting direct transcriptional control of β7 expression by the CB_2_R-RA axis^47^. However, *CNR2* knockdown had no effect on RA-induced integrin β7 expression, indicating that RA does not rely exclusively on CB_2_R to mediate β7 induction. This increased α4β7 expression has functional implications too as demonstrated by adherence assays using immobilised MAdCAM-1 and endothelial monolayers. Once again CB_2_R agonism increased adherence while inverse agonism or knockout suppressed it. This is in contrast with previous reports of leukocytes CB_2_R activation leading decreased adhesion to brain endothelium and migration across the blood-brain barrier^43^. However, that disparity likely reflects the driving of gut-specific integrin expression instead.

As a G-protein coupled receptor, CB_2_R primarily couple to G_o_ proteins leading to inhibition of adenylate cyclase, cyclic adenosine monophosphate (cAMP), and calcium channels^4^. This then triggers activation of mitogen-activated protein kinases (MAPKs), including p38, Jun-terminal kinase, extracellular signal-regulated kinase (ERK)1/2, and p44/42 MAPK^5–7^. Transcription factor analysis of the involved promoter sequence revealed a predominance of binding sites for factors downstream of the Ras/Raf/MEK/ERK pathway, notably p53 and c-Jun. Multiple Ras/Raf/MEK/ERK-associated transcription factors are predicted to bind regulatory regions critical for CB_2_R-mediated α4β7 integrin expression in T cells, further implicating this signalling axis. While Raf/MEK/ERK activation drives integrin α6 and β3 expression, the contribution to integrin β7 has not been previously described^48^. Using a pan-Raf inhibitor, we were able to confirm the requirement for this pathway in CB_2_R-mediated α4β7 upregulation. To clarify this mechanism, we focused on ERK signalling, as it acts downstream of CB_2_R activation and emerged as a key regulator of CB_2_R-driven transcriptional responses in T cells during our pathway analysis. Signalling through ERK is also essential for the expression of integrins such as α2b1, αvβ3, and αvβ5^49^. RA signalling drives ERK activation in T cells, as evidenced by increased phosphorylation of both ERK1 and ERK2^50^. CB_2_R signalling also activates ERK phosphorylation in both neuronal and immune cells^33, 51^. Given the established role of ERK signalling in integrin regulation and its activation by both RA and CB_2_R pathways, we hypothesized that ERK mediates CB_2_R-driven α4β7 expression in T cells. Using a time-course flow cytometry assay, CB_2_R activation significantly increased ERK1/2 phosphorylation at T202/Y204, supporting this mechanism. These results indicate that indeed Ras/Raf/MEK/ERK cascade is a key driver of CB_2_R-induced α4β7 expression.

Previous studies indicate that phytocannabinoid treatment promoted CD4^+^ cell gut-homing in a non-human primate SIV model^29^. If the same happens in humans, this may go some way to explaining how chronic cannabis use activating CB_2_R might lead to worsening disease outcomes for IBD patients^27^. CB_2_R expression is upregulated in immune cells across multiple chemically induced colitis models (mustard oil, TNBS, DSS)^14, 15^. Cnr2 is also significantly upregulated in the inflamed ileum of TNF^ΔARE^ mice, with corresponding increases in CB_2_R protein levels compared to WT^12^. Elevated CB_2_R expression in ileal and colonic biopsy samples from CD and UC patients, respectively, mirrors findings from animal models^16–18^. Numerous studies have also indicated CB_2_R to be overexpressed on CD4^+^ T cells isolated from human IBD patients^15, 18, 52, 53^. These studies highlight the potential role of this receptor in IBD progression and intestinal immune regulation.

Antibody blocking α4β7 with Vedolizumab selectively inhibits the intestinal trafficking of α4β7-expressing T cells, with 41.8% and 39% clinical remissions rate in both UC and CD patients, respectively. Therefore, the finding that CB_2_R signalling regulates the expression of α4β7 on T cells offers therapeutic potential for IBD. Several studies have previously described attenuated colitis following CB_2_R activation in murine models in contrast to our findings^54^. Notably, our results align more closely with clinical data from human IBD patients, which show no benefit and, in some cases, worsening outcomes with cannabinoid exposure^27, 55, 56^. We propose three potential explanations for this discrepancy. Firstly, the effect may be site-specific as our adoptive transfers studies saw no difference in T cell trafficking to the colon which might be important for colitis studies, only the ileum which was then reflected in our ileitis model. Secondly, our model employs chronic ileitis in 20-week-old mice, better mimicking the long-term nature of human IBD compared to the acute colitis studies. Finally, the TNF^ΔARE/+^ model, like human disease, is highly T cell-dependent as unfractionated CD4^+^ T cells are sufficient to transfer disease in lymphopenic recipients^41^. CD4⁺ T cell-specific CB_2_R knockout allows for targeted interrogation of T cell-mediated mechanisms in this model. In contrast, acute colitis models remain disease-prone even in lymphopenic settings consistent with a decreased role for T cells in those models^57^. Together, our findings highlight the need to better understand molecular mechanisms driving CB_2_R-induced immune responses in IBD and pave the way for future development of novel CB_2_R-targeting drugs to broaden the arsenal in the ongoing battle against this incurable lifelong disease. This study also provides mechanistic insight underpinning the known risks associated with cannabis use among people with IBD.

---

**Figure S1.** IPA provides insights into the molecular pathways implicated following CB_2_R blockade in T cells. (A) Pathway analysis indicating the molecular pathways downregulated following CB_2_R blockade. Node colour is indicative of the degree of regulation. Node size is based on the number of genes involved in that given pathway. (**B**) Top gene-gene interaction network constructed using IPA based on the genes specifically regulated following CB_2_R blockade. Node colour is indicative of the direction and degree of regulation, with blue indicating downregulation and red indicating upregulation.

**Figure S2.** IPA provides insights into the molecular pathways implicated following CB_2_R activation in T cells. (B) Pathway analysis indicating the molecular pathways upregulated following CB_2_R activation. Node colour is indicative of the degree of regulation. Node size is based on the number of genes involved in that given pathway. (**B**) Top gene-gene interaction network constructed using IPA based on the genes specifically regulated following CB_2_R activation. Node colour is indicative of the direction and degree of regulation, with blue indicating downregulation and red indicating upregulation.

