## Supplementary figures and images for "Cannabinoid CB2R regulates T cell gut-homing in preclinical Crohn’s model"

### Supplemental Figures 1 & 2

#### Figure S1 Pathway analysis

A.

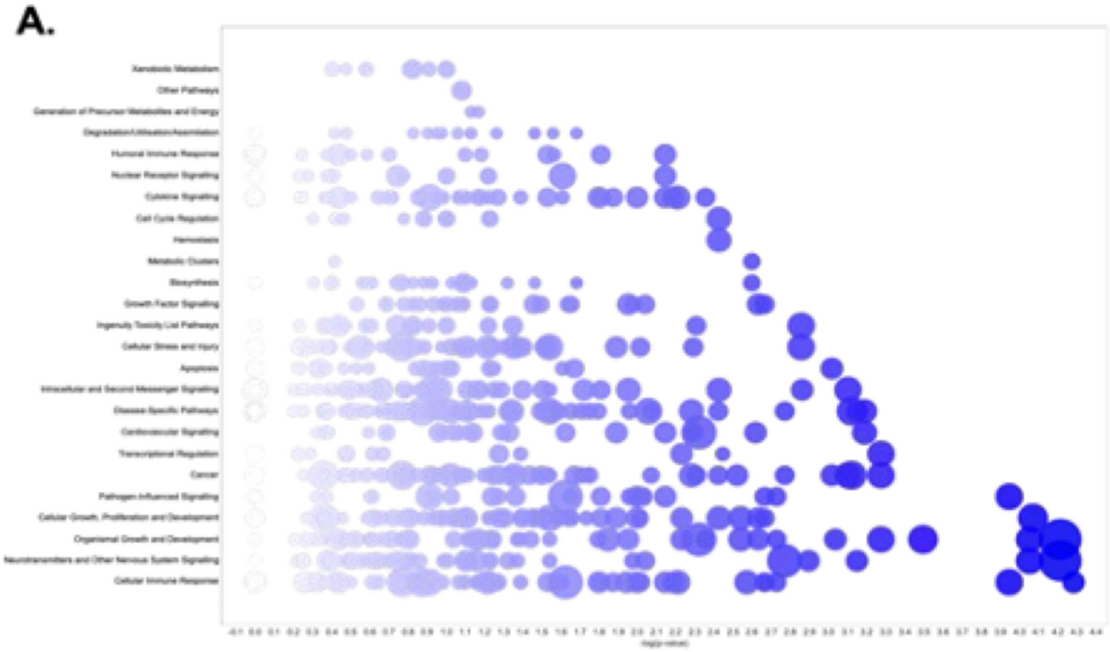


B.

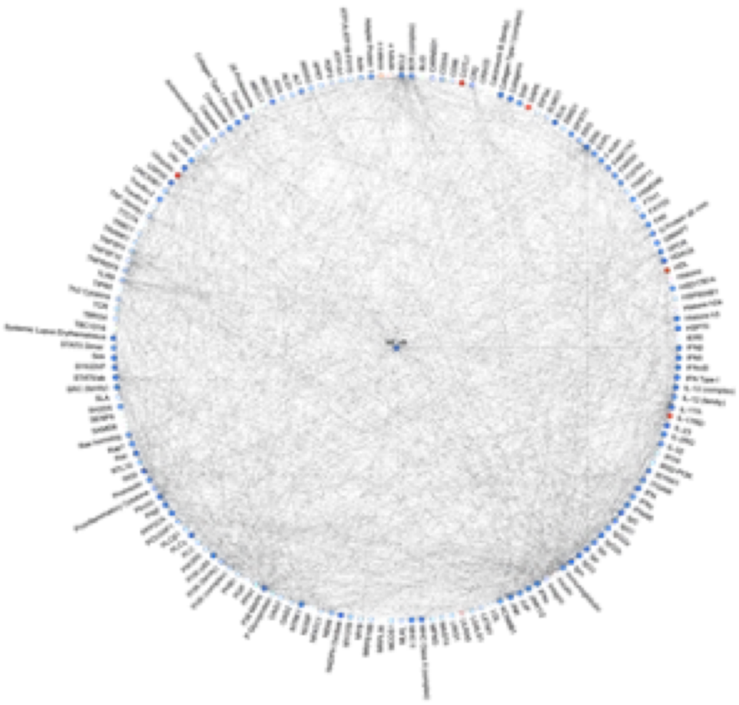

#### Figure S2 Pathway analysis

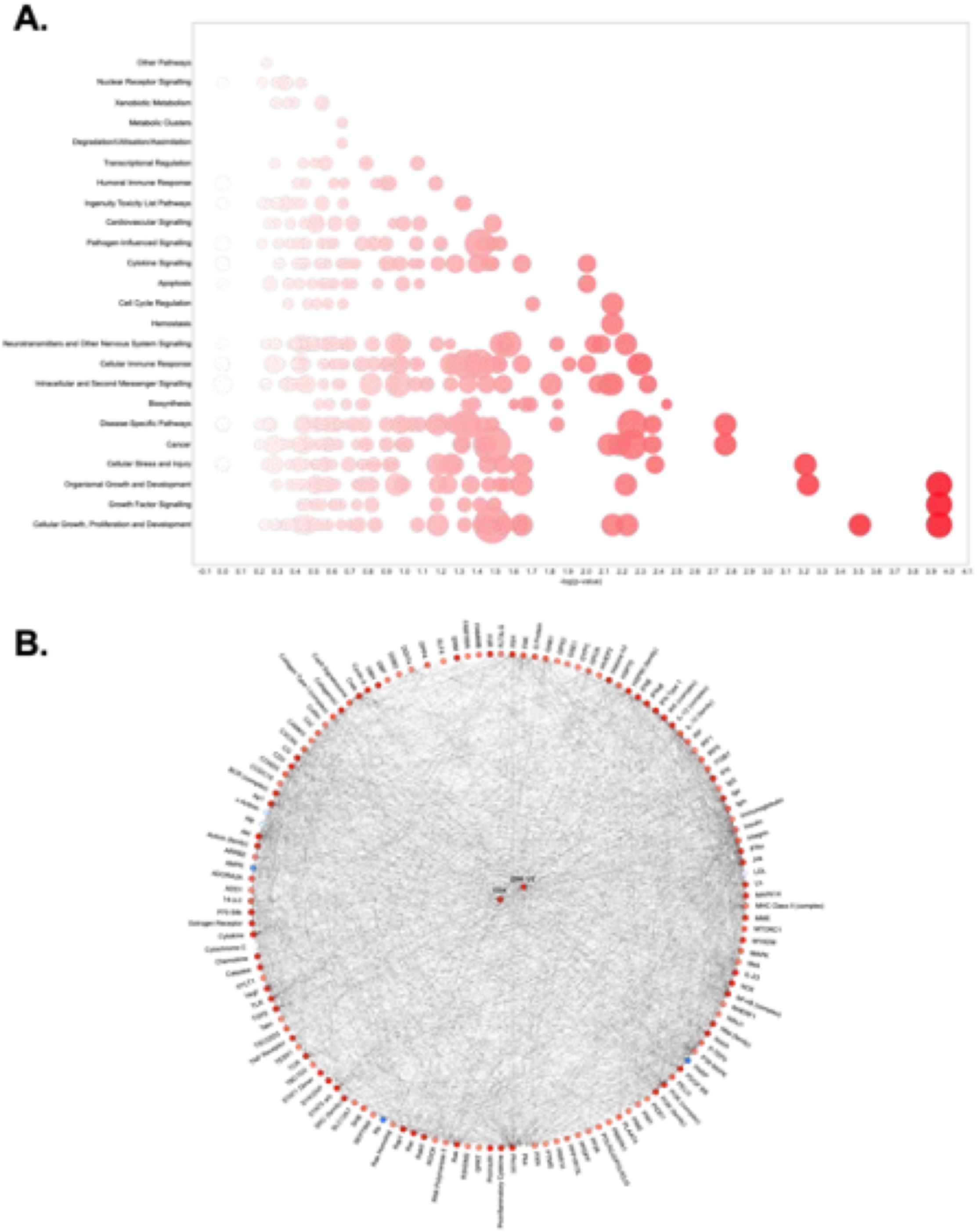
